# Epigenetic signatures of chronic social stress in stress-susceptible animals

**DOI:** 10.1101/690826

**Authors:** Nicholas O’Toole, Tie-Yuan Zhang, Xianglan Wen, Josie Diorio, Patricia P. Silveira, Benoit Labonté, Eric J. Nestler, Michael J. Meaney

## Abstract

Exposure of mice to chronic social defeat stress (CSDS) produces depressive- and anxiety-like behaviors and widespread transcriptomic changes in several brain regions in susceptible animals. Here we present the first study of genome-wide cytosine methylation patterns of mice susceptible to CSDS using whole-genome bisulfite sequencing on DNA from the nucleus accumbens, a critical region for CSDS effects on behavior. We found extensive evidence for differential methylation following exposure to CSDS in susceptible animals, with a greater proportion of CG hypermethylation than hypomethylation in CSDS-susceptible mice compared to non-stressed controls; non-CG methylation shows the opposite trend. Several genes previously implicated in the effects of CSDS are among those with the greatest number of differentially methylated sites, including estrogen receptor alpha (*Esr1*), the deleted in colorectal cancer (*Dcc*) gene and *Cacna1c*, which has been associated with a range of psychiatric conditions. Informatic analysis of DM sites revealed a gene network with ß-catenin as the hub gene of a network that included the ß-catenin-related WNT/frizzled signaling pathway as well as both *Esr1* and *Dcc*. Finally, we found considerable overlap between DM genes associated with CSDS in susceptible animals and those associated with human neuroticism in a genome-wide association study. Analysis of these overlapping genes revealed ‘WNT signaling’ as the top pathway, which features ß-catenin as the primary hub gene. These findings reveal a striking convergence between the molecular pathways identified through either transcriptional or epigenomic analyses of the mouse model of susceptibility to chronic stress and the genomic architecture of increased stress susceptibility reflected in neuroticism in humans.

## Introduction

Prospective studies confirm that exposure to stress together with enhanced stress reactivity predicts a greater risk for later depression ^1^. Likewise, neuroticism, which is in part defined by increased stress reactivity, is an endophenotype closely associated with depression and anxiety disorders ^2^. Neuroticism influences the interaction between stressful events and negative affect, including symptoms of depression and anxiety, and may thus be considered as reflecting greater susceptibility to stress. Nagel et al. (2018) used a large GWAS meta-analysis that implicated 599 genes for which variants were significantly associated with neuroticism ^3^. Subsequent informatic analyses revealed enrichment for specific cell types, including ‘dopaminergic neuroblasts’ and ‘striatal medium spiny neurons’ ^3^. Pathway analyses using this gene set identified the ‘behavioral response to cocaine’ which involves the mesocorticolimbic dopamine reward circuit. This circuit is also closely associated with stress-induced behavioral effects, including anxiety- and depressive-like behaviors ^4^.

There is compelling evidence for the role of the mesocorticolimbic dopamine circuit in mediating individual differences in stress reactivity in studies of chronic stress with model systems ^5^. Optogenetic studies show that multiple corticolimbic structures converging on the nucleus accumbens (nAcc) are involved in the activation of behavioral stress responses, including the medial prefrontal cortex, the ventral hippocampus and the ventral tegmental area ^6–8^. Activity within these regions and in particular in pathways that project to the nAcc distinguish animals that show susceptibility to chronic stress as defined by a number of behavioral measures of depression- and anxiety-like states ^9^.

Genome-wide transcriptional analyses also implicate the nAcc as a critical site for stress-induced behavioral alterations and distinguish animals that are susceptible to the effects of chronic social stress ^7^. These transcriptomic findings suggest underlying epigenetic mechanisms associated with exposure to chronic stress in susceptible animals ^10^. Here, we sought to define the epigenetic landscape associated with exposure to chronic social defeat stress (CSDS), a model commonly used to explore individual differences in susceptibility to chronic stress and with strong ethological validity. We focused on DNA methylation, which is a highly stable epigenetic modification and associated with the capacity for regulating transcriptional activity ^11–14^. Previous studies using this model reveal that a substantial portion of animals exposed to CSDS are resilient (i.e., behaviorally indistinguishable from non-stressed controls). Such marked individual differences in susceptibility complicate research designs attempting to define the molecular pathways that underlie the pathophysiological consequences of exposure to chronic stress. To circumvent this limitation, we first confirmed the susceptibility of CSDS-exposed animals in this epigenetic analysis with behavioral testing. This approach is essential in defining the epigenetic modifications associated with exposure to CSDS related to the induction of depressive- or anxiety-like behaviors. We then performed whole-genome bisulfite sequencing (WGBS) of genomic DNA samples derived from the nAcc (core and shell regions) from individual CSDS-susceptible and non-stressed control mice to define variably methylated regions at single-base pair resolution genome-wide including the characterization of CSDS-related changes in both CpG and non-CpG (i.e., CHG and CHH) methylation. The latter is of particular relevance considering the unique prevalence of this epigenetic mark in brain ^15^. Our findings reveal widespread differences in both CG and non-CG methylation with a marked alignment with previous transcriptional analyses using the CSDS model. Informatic analyses identify *Esr1* as a candidate hub for a network of genes showing differential methylation, particularly at non-CG sites, between CSDS-susceptible and control animals, consistent with previous studies showing that *Esr1* overexpression in the nAcc regulates susceptibility to CSDS ^16^. This analysis also reveals enrichment for differentially-methylated genes converging on gene networks that associated with canonical WNT/ß-catenin signaling. These findings suggest that transcriptional pathways that underlie the effects of chronic stress in susceptible animals are associated with epigenetic profiles involving DNA methylation in genes implicated in neurogenesis and synaptic plasticity. Finally, we explored the clinical relevance of these data sets by comparing the list of differentially-methylated genes in CSDS-susceptible mice with those associated with neuroticism in the Nagel et al. (2018) GWAS ^3^. We found significant overlap, and informatic analyses focused on the overlapping targets again identified the WNT/ß-catenin signaling as the primary candidate pathway.

## Results

### Mapping of bisulfite sequencing reads

Independent genomic DNA samples from the nAcc of 5 control and 4 CSDS-susceptible mice (Fig. 1a) were bisulfite treated and the prepared libraries were processed in Illumina HiSeq X at a sequencing depth of 30 x genome coverage for six technical replicates per sample. The number of sequencing reads for all 54 FASTQ files ranged from 71.1 to 130.0 million. Bismark mapping to the mouse mm10 genome after the pre-processing steps (see Methods) resulted in mapping percentages ranging from 72.7% to 76.5%. The non-conversion rate was calculated by spiked-in unmethylated phage lambda DNA and found to be no more than 0.1%. Bismark BAM files from the six technical replicates for each sample were sorted and merged for input to the methylKit package resulting in 9 BAM files each containing information on over 400 million mapped sequencing reads. MethylKit was applied to generate methylation call files for each CG or non-CG in the genome using a minimum coverage of 10 reads per position. Raw sequencing read files and methylation call files have been submitted to the Gene Expression Omnibus (GEO) under accession GSE126955. Approximately 75% of cytosines in CG context, and 2.3% in non-CG contexts showed some degree of methylation across all samples, with no significant differences in total methylation between the control and CSDS-exposed susceptible groups. These genome-wide levels of the various forms of methylation are consistent with previous studies using WGBS on mouse brain ^15,17,18^.

**Figure 1.**
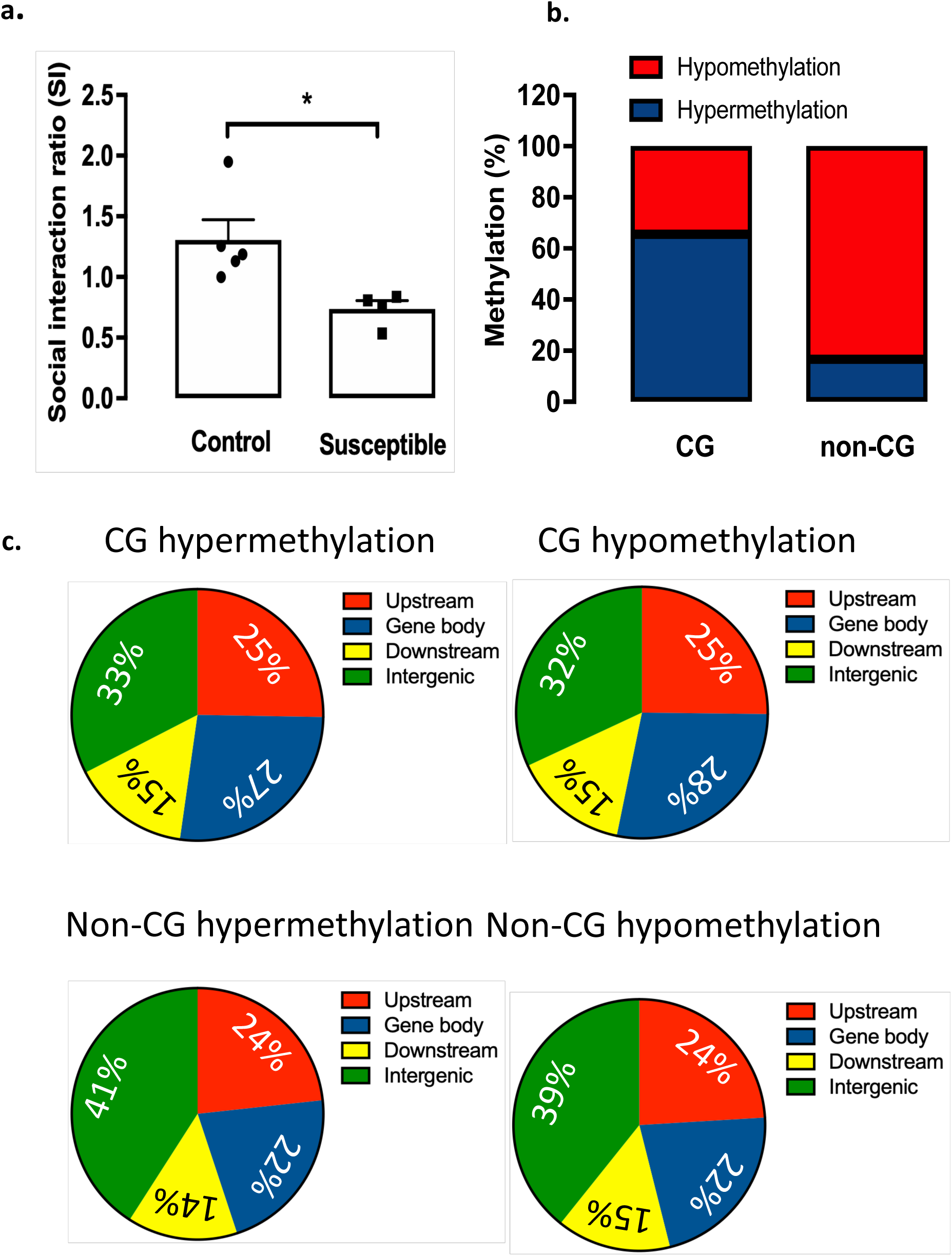
Social defeat and genome-wide DNA methylation. **a)** Mean +SEM of social interaction (SI) ratio of CSDS-susceptible and non-CSDS exposed control mice (t _(7)_= 2.86, * p = 0.024). **b)** The proportion of hypermethylated bases (CSDS > controls) is greater than that of hypomethylated bases at CG sites. Non-CG sites show greater hypomethylated bases than hypermethylated bases. **c)** Pie charts of differentially methylated cytosine bases in sequence context.

### Differentially methylated sites in nAcc between controls and CSDS-susceptible mice

The default methylKit significance cut-offs of corrected p-value (q) < 0.01 and absolute value of methylation percentage difference (|Δβ|) > 25% were used to identify 79,507 differentially-methylated (DM) CG sites between control and CSDS-susceptible mice. Of these sites, 51,915 showed hypermethylation in samples from CSDS-susceptible over control animals and 27,592 hypomethylation (Fig. 1b). “Hypermethylation” is defined as significantly greater levels of methylation in CSDS-susceptible samples over controls for a given cytosine; “hypomethylation” reflects less methylation in CSDS-susceptible samples then controls.

DM sites were classified by their distance from transcription start sites (TSS) and end sites (TES) of annotated Ensembl genes. Coordinates further than 20 kilobases (kb) upstream from a TSS or 20 kbs downstream from a TES were classified as “intergenic”. Those sites closer to the TSS or TES were classified as “upstream” or “downstream” respectively. DM sites between the TSS and TES were classified as “gene body”. The 20 kb cut-off is somewhat arbitrary but will certainly contain up-or down-stream regulatory elements for the nearest gene other than more distant elements that reflect chromatin looping. A majority of DM sites are in or near annotated genes and there are more DM CGs upstream of genes than downstream. Separate pie charts for hyper- and hypo-methylated CG and non-CG sites (Fig. 1c), show highly similar distributions of gene location classifications, which is consistent with previous findings on the proximal relation between CG and non-CG methylation ^15,17^.

Differential methylation statistics were also computed for all cytosines in a CHG or CHH context (i.e., non-CG methylation where “H” represents an A, T or C nucleotide) with a minimum coverage of 10 sequencing reads, with numbers of differentially methylated bases using the q < 0.01 Δβ > 25% cut-offs. We find a greater proportion of hypermethylated bases in CG context compared to non-CG context (Fig. 1b) and discuss this finding below. The differences in the proportions of hyper- and hypomethylated bases for CG and non-CG contexts is significant (P < 1e-20, two-proportion z test). As was the case for CGs, the relative proportions of hyper- and hypo-methylated bases in the non-CG pie charts are indistinguishable from each other by eye (Fig. 1c).

As noted above, there is a striking bias in the methylation state of DM sites in nAcc that distinguish control and CSDS-susceptible animals (Fig. 1b). Non-CG sites in DM regions are significantly (two-proportion z test: P < 1e-22) more likely to be hypo-compared to hyper-methylated. The opposite is the case for CG methylation. Interestingly, a similar bias towards hypomethylated DM regions is observed in the comparison of non-CG methylation levels in induced pluripotent stem cells compared with embryonic stem cells ^19^. The very distinct differences in the methylation levels of CG and non-CG in the DM regions (hyper-vs hypo-methylation) suggests different causal mechanisms for the formation of CSDS-related differential CG and non-CG methylation (i.e., differences in non-CG methylation are not merely a stochastic by-product of a common process). Non-CG methylation, unlike CG methylation, is largely asymmetric with evidence for distinct pathways. Non-CG methylation is also enriched in neurons as compared to glia, as distinct from CG methylation patterns. Non-CG methylation can be sustained in the absence of DNA methyltransferase 1 (DNMT1) and appears more closely associated with the *de novo* activity of DNMT3a/b as well as DNMT3-like protein (DNMT3l) ^20–24^. Moreover, non-CG methylation tends to accumulate in regions with higher levels of chromatin accessibility ^15,24^, suggesting some degree of independence from CG methylation, which associates with reduced chromatin accessibility. Nevertheless, increased levels of non-CG methylation in promoter regions, like that of CG methylation, associate with decreased transcriptional activity ^15,24^, which suggests a common influence on transcription. Dynamic periods of non-CG remodeling associate with peak developmental periods of both synaptogenesis and synaptic pruning in human and mouse brain and coincides with enhanced DNMT3a expression ^15^. Interestingly, CA methylation, which is the predominant form of non-CG methylation, is the form of DNA methylation most affected by peripubertal environmental enrichment ^18^, a condition that likewise associates with synaptic remodeling. These findings are interesting to consider in light of the candidate functions emerging from our subsequent informatic analyses (see below) that highlight pathways closely associated with neurogenesis and synaptic plasticity.

### Differential methylation of genes associated with susceptibility to chronic stress

A total of 9,901 genes contained at least one DM site within the gene body, representing approximately 18% of annotated mouse genes from Ensembl. We used a relatively conservative approach and restricted our analyses to those regions showing more widespread differential methylation. Murine genes containing at least 7 DM sites within the gene body and at least 6 DM sites within 20 kb upstream of the TSS were identified for further investigation. Despite the rather conservative criteria, there were nevertheless 626 such genes, representing a little over 1% of the annotated genes in mouse. These regions are referred to as “DM genes” on the basis of the relatively large number of differentially-methylated cytosine bases within or upstream of the genes. Data for these genes and the DM sites within or in close proximity to them are contained in Supplementary Table 1.

Interestingly, several of the most highly differentially methylated genes (Fig. 2a-c) have been identified in previous studies of transcriptional changes associated with behavioral responses to CSDS. Notable among these is *Esr1*, which encodes estrogen receptor alpha. Our previous studies ^16^ show that overexpression of *Esr1* selectively within the nAcc significantly reduces susceptibility to CSDS. That same study highlights 31 differentially expressed genes between susceptible and resilient mice known to interact with *Esr1* ^16^. *Three of these genes are found in the present DM gene list, Gabbr2, Adcy1* and *Sulf1*, reflecting a significant overlap (P < 0.045, Exact Hypergeometric test).

**Figure 2.**
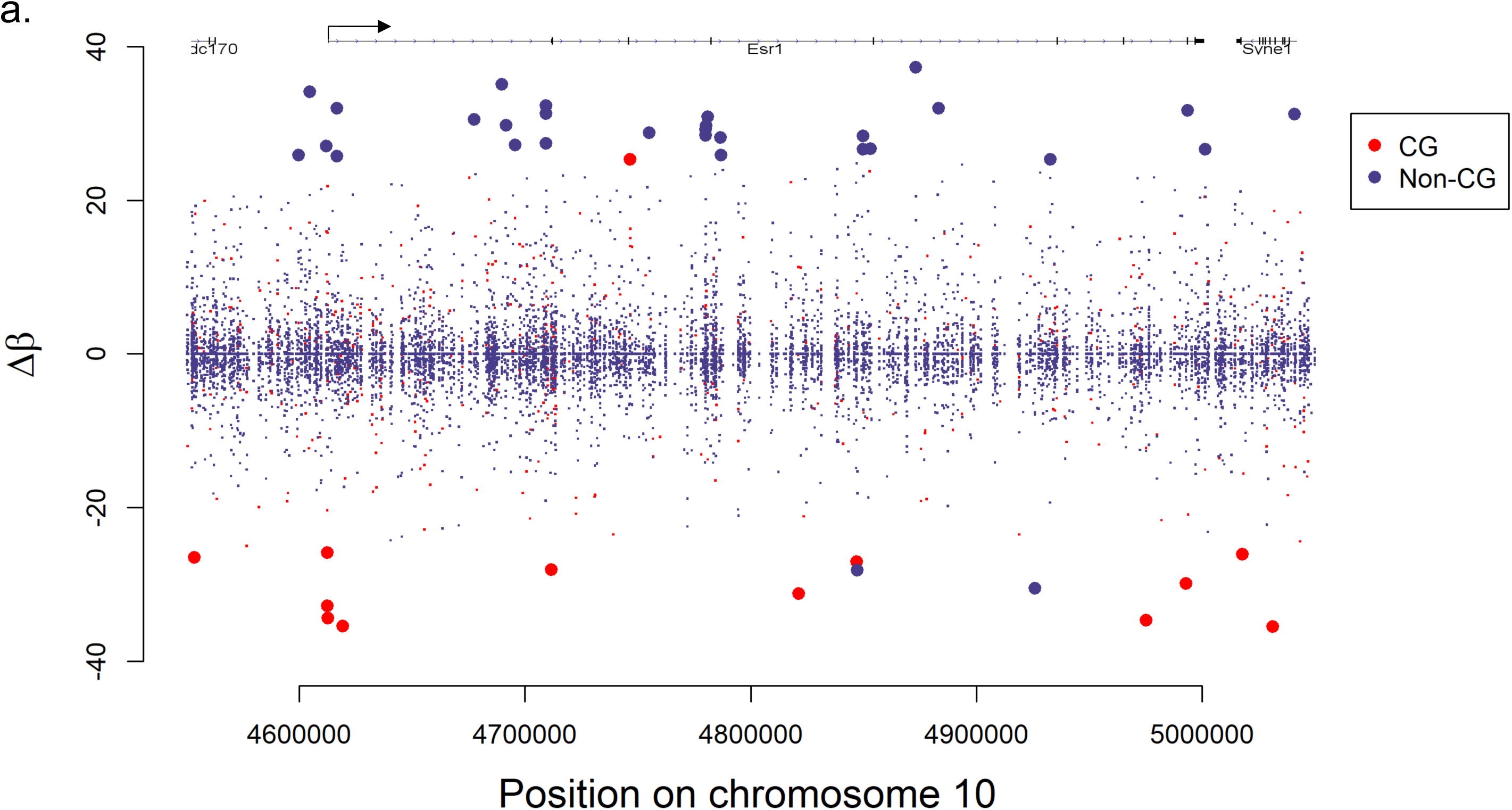
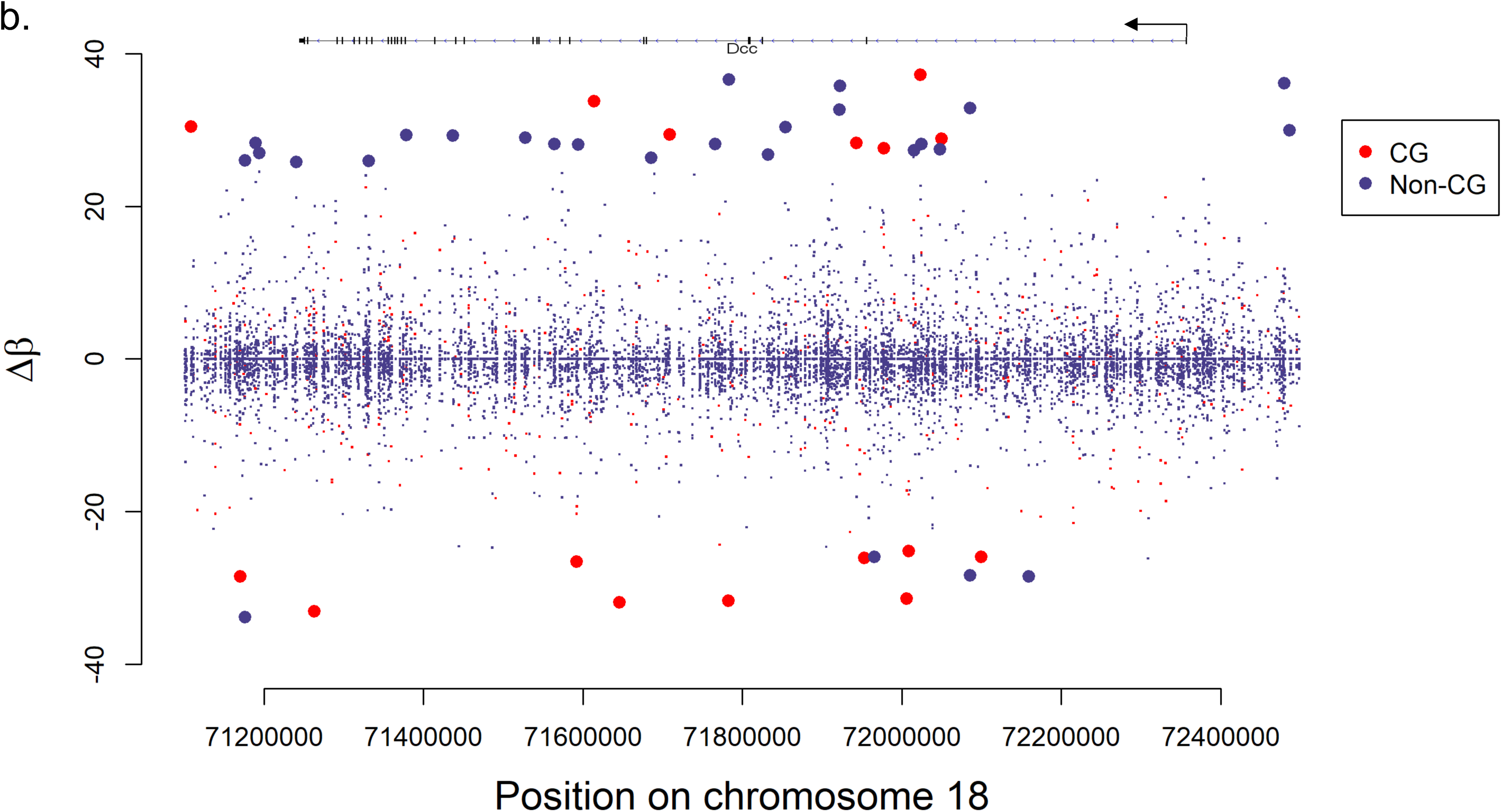
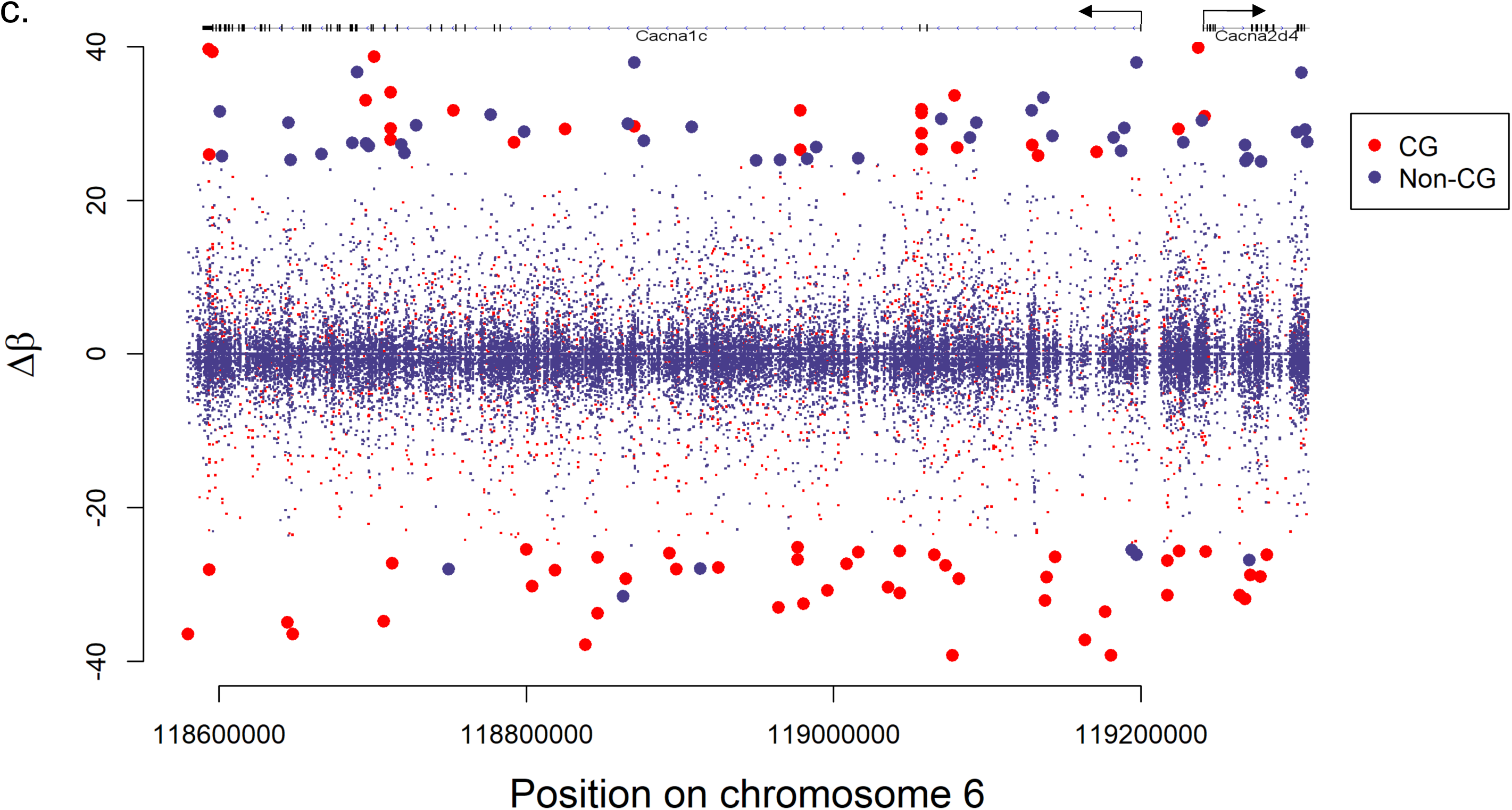
Methylation differences of cytosines at genes of interest. **a)** *Esr1*, **b)** *Dcc*, **c)** *Cacna1c*. Larger sized points represent differentially-methylated bases satisfying the significance cut-offs of |Δβ|) > 25% were used to identify 79,507 differentially-methylated (DM) CG Δβ|Δβ|) > 25% were used to identify 79,507 differentially-methylated (DM) CG > 25 and q < 0.01. Bases hypermethylated in susceptible samples over controls have negative Δβ values; positive values correspond to hypomethylation.

The DM gene list also includes the *Dcc* gene. *Dcc* transcription in the prefrontal cortex (PFC) is associated with susceptibility to CSDS in mice and with major depressive disorder in humans ^25^. Although these results were obtained in the PFC, Bagot et al. (2016) show a significant correlation between transcriptional changes and overlap of differentially expressed genes in the nAcc and PFC in response to CSDS (Fig. 2a, c of Bagot et al. 2016)^7^, significantly more than that occurring between any other brain region examined. Torres-Berrio et al. also show that the microRNA miR-218 regulates the transcription of *Dcc* ^25^. We find no DM CGs within 20 kb of mir-218a, but there were two hypermethylated CGs approximately 2 kb downstream of miR-218b.

*The Cacna1c* gene showed a remarkable level of differential methylation between control and CSDS-susceptible mice with 38 DM sites within the gene body (Fig. 2c). Terrillion et al. (2017) show in mouse nAcc that decreased expression of the *Cacna1c* gene, which codes for the α1C subunit of the Ca_v_1.2 L-type calcium channel subunit (LTCC), is associated with susceptibility to social stress ^26,27^. Furthermore, the neighboring gene, *Cacna2d4*, as well as other LTCC subunits, including *Cacna1a, Cacna1e, Cacna1i* and *Cacna2d3*, are all contained in the DM gene list of the current study. GWASs in humans identify significant association between polymorphisms with in the *Cacna1c* gene and a range of psychiatric disorders ^26,27^ including bipolar disorder ^28^ and depression ^29–31^; hypermethylation in the gene associates with bipolar disorder ^32^.

The overall finding that DM non-CG sites are more likely to be hypo-rather than hyper-methylated, with an opposite trend for CG methylation, is also apparent in the genes noted above. Fig. 2a-c shows that a large majority of the DM non-CG bases in or near *Esr1, Dcc* and *Cacna1c*, indicated by large blue points, are hypomethylated (Δβ > 0). In contrast, DM CGs in these genes are generally hypermethylated.

Finally, we used previously published data sets to explore the relation between our DM genes and those showing differences in transcription as a function of susceptibility to CSDS. Bagot et al. (2016) used RNA-seq to identify 358 genes differentially in the nAcc between control and CSDS-susceptible mice ^7^. We used a cut-off of at least 4 DM CGs in the gene body to define DM genes and examined the overlap with the differentially expressed genes identified in the Bagot et al. study, which revealed 34 overlapping genes (Hypergeometric test: P < 0.005).

### Gene network analyses

We mapped DM regions to genes and used Metacore^®^ (Clarivate Analytics) to identify gene ontologies and networks (Fig. 3). We performed separate analyses for DM genes located in the upstream and gene body regions based on the known DNA methylation profiles of actively transcribed and silenced genes ^13,33^. The former is characterized by hypomethylation in the upstream regulatory regions and hypermethylation in the gene body; the reverse is the case for transcriptionally silenced genes.

**Figure 3.**
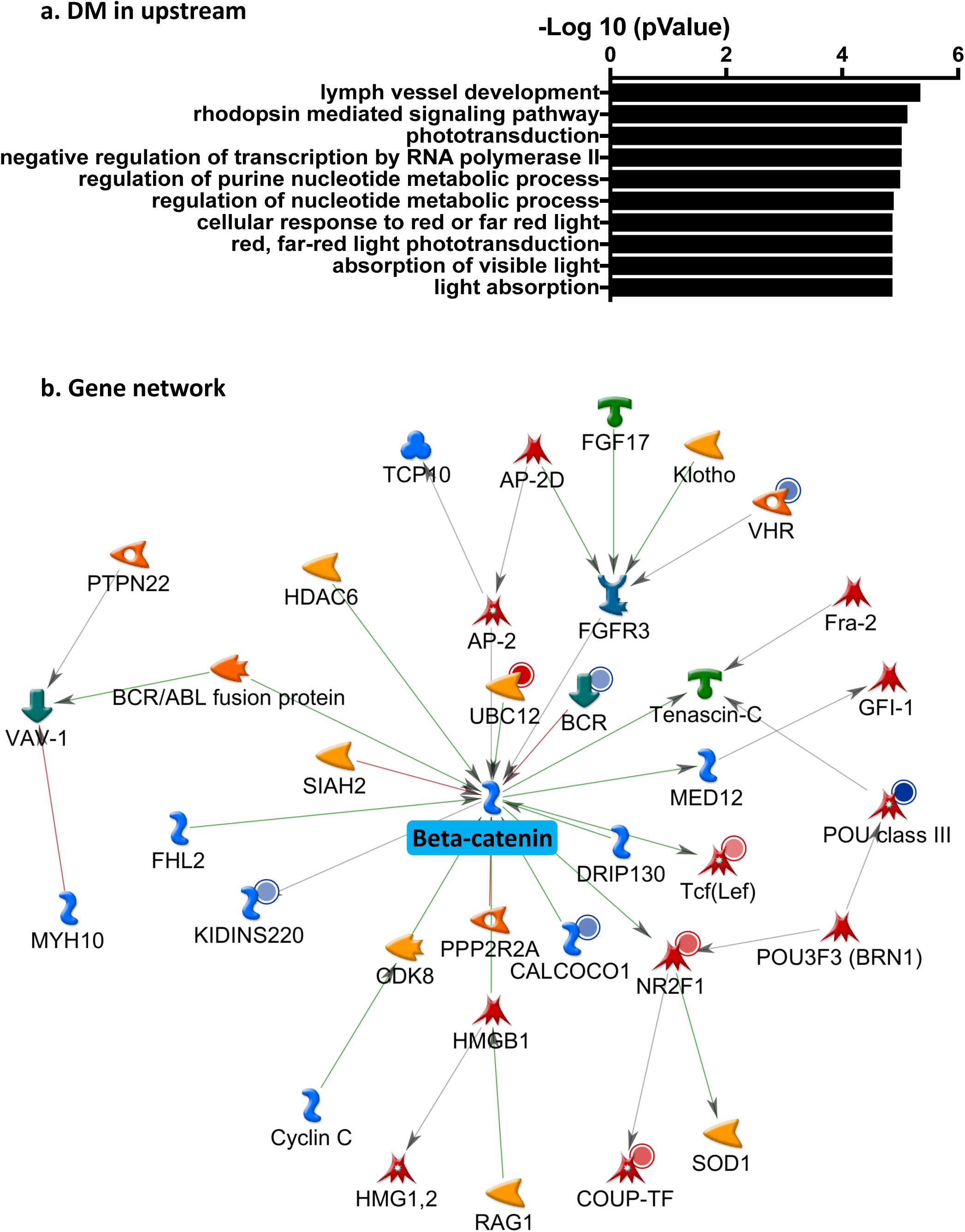
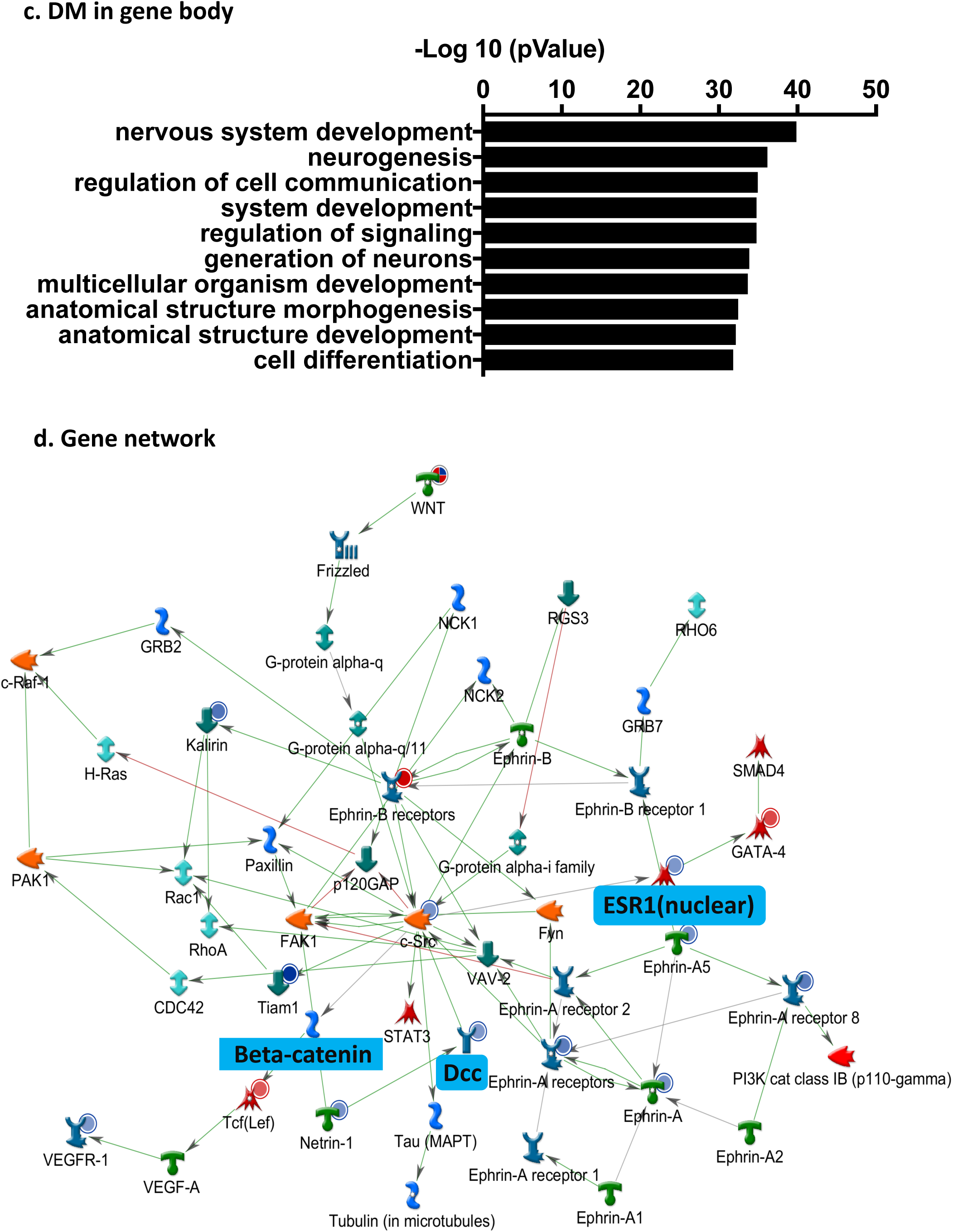
Epigenetic network signature of CSDS in susceptible mice. **a**) GO analysis of genes differentially methylated at upstream sites between CSDS-exposed susceptible and control mice. **b**) Analysis of genes differentially-methylated in the upstream region between CSDS-susceptible and control mice reveals a top network with ß-catenin as the hub gene. **c)** GO analysis of genes differentially methylated in the gene body between CSDS-exposed susceptible and control mice. **d)** Analysis of genes differentially methylated in the gene body region between CSDS-exposed susceptible and control mice reveals a top network that includes Ephrin receptors and their ligands as well as ß-catenin, Dcc and netrin, and Esr1. The graded intensity of blue dots represents increased DNA methylation in the genes in CSDS-exposed susceptible compared to control mice. The red dots represent decreased DNA methylation in CSDS-exposed susceptible compared to control mice.

Analysis of upstream DM regions revealed a gene network with ß-catenin as the hub gene (Fig. 3 a,b). This gene has been strongly implicated as a key determinant within the nAcc of susceptibility to chronic stress ^4^. Mice with a local knockout of ß-catenin or overexpressing a ß-catenin dominant negative mutant in nAcc show enhanced susceptibility to CSDS, while selective overexpression of ß - catenin in nAcc principal neurons enhances resilience. The expression level of ß-catenin is up-regulated in hippocampus by several antidepressant medications ^34^. Likewise, analysis of DM regions within gene bodies in nAcc in our study identified a network that included the ß-catenin-related WNT/frizzled signaling pathway (Fig. 3c.d), which operates in conjunction with ß-catenin to regulate neuroplasticity ^35^. Importantly, this network included two targets that showed the highest level of differential methylation, notably *Esr1* and *Dcc*. This network also includes a number of ephrin receptors known to be associated with WNT and ß-catenin signaling ^35–37^.

### DM genes in CSDS-susceptible animals overlap with neuroticism-associated genes and identify common molecular pathways

Neuroticism is a stable trait that involves enhanced susceptibility to stress and a risk for depression and anxiety disorders ^2^. There is significant genetic co-morbidity between neuroticism and both anxiety and depression^38^. Nagel et al. (2018) reported 599 genes in which variants were significantly associated with neuroticism in a large GWAS meta-analysis ^3^. We compared these genes to our list of genes associated with DM regions both upstream and in the gene body region using Metacore. We found 30 common genes, with 25/30 overlapping with our gene body list including *Esr1, Dcc* and *WNT3*. The enrichment analysis shows the significant GO term enrichment for ‘cell development’ (p>10^-15^). We then performed an enrichment analysis on the genes overlapping between the gene body DM list in the mouse analyses and the genes identified in GWAS for neuroticism (Fig. 4, Supplementary Figure S1a. b.). The most highly significant pathway was that of ‘WNT signaling’ (p<10^-6^) and the resulting map for this pathway clearly reveals ß-catenin as the primary hub for the WNT signaling pathway. These findings reveal a striking convergence between the molecular pathways identified through either transcriptional or epigenomic analyses of the mouse model of susceptibility to chronic stress and the genomic architecture of increased stress susceptibility reflected in neuroticism in humans.

**Figure 4.**
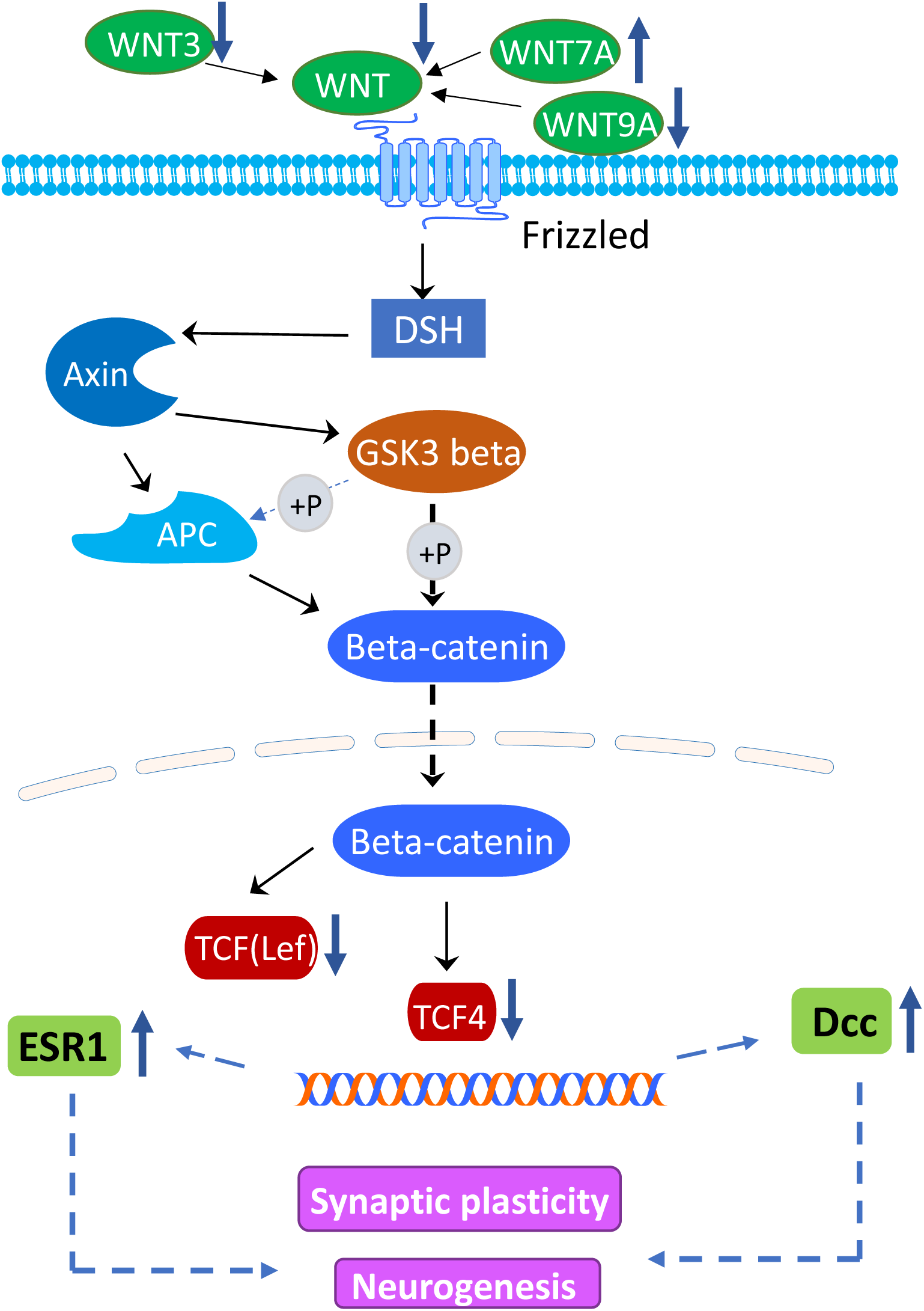
Illustration of the common gene pathways between differentially methylated genes in gene body of CSDS-exposed susceptible and control mice and those genes identified in the GWAS for neuroticism. Enrichment analysis of the overlapping genes reveals the WNT-ß-catenin pathway as the top biological process that linked to neurogenesis and synaptic plasticity. The blue arrows represents differentially-methylated genes in the CSDS-exposed susceptible and control mice.

## Discussion

We used WGBS to define cytosine methylation at single nucleotide resolution in the nAcc of mice behaviorally-defined as CSDS-susceptible compared with non-stressed controls. This study represents what is to our knowledge a novel genome-wide analysis at single-base pair resolution of DNA methylation profiles associated with stress susceptibility. Despite relatively conservative criteria, our results reveal widespread differences in multiple forms of cytosine methylation. We found striking divergence in the profiles of the different forms of DNA methylation in the nAcc, with hypermethylation at CG sites in CSDS-susceptible compared with control animals coupled with decreased levels of non-CG sites. Hing et al (2018) reported broad changes in DNA methylation following exposure to CSDS using a methyl-capture sequencing with samples from the dentate gyrus ^39^. This approach does not afford single-base pair resolution and the identification of non-CG forms of methylation, which, together with tissue differences, precludes a direct comparison with our data set. However, together the two studies support the idea that exposure to chronic stress associates with widespread alterations to the methylome.

Interestingly, the regional distribution of CG and non-CG sites across the genome is highly similar (Fig. 1c). Moreover, non-CG sites tend to be found in proximity to those bearing CG marks. These findings suggest that exposure to CSDS in susceptible animals initiates the conversion of non-CG to CG sites. CSDS associates with increased expression of DNMT3a in the nAcc ^40,41^. DNMT3a binding regions are enriched for non-CG methylation ^15^, and DNMT3a complexed with DNMT3l, which functions as a regulatory factor and produces *de novo* CG methylation ^42^. CSDS-induced changes in expression levels of multiple DNMTs might catalyze the conversion of non-CG to CG sites. We must caution that, given the present gap in our knowledge of the mechanisms for the distinct formation of CG and non-CG sites, this remains a matter of speculation. However, this hypothesis is consistent with the findings that exposure to chronic stress increases DNMT3a expression in multiple brain regions, including the nAcc ^40,41^ and mediates behavioral effects ^40,41^. Overexpression of DNMT3a in the nAcc enhances susceptibility to CSDS in male mice ^43^. Interestingly, *Dnmt3a* expression levels are increased in the nAcc of depressed humans ^43^.

Informatic analyses were conducted selectively based on differential methylation in upstream and gene body regions based on the region-specific relationship of methylation with gene transcription. We found impressive convergence of the gene networks emerging from these independent analyses on the WNT/ß-catenin signaling pathway (Fig. 3b. d). WNT ligands act through frizzled receptors to activate β-catenin. The activation of this canonical WNT signalling cascade results in the stabilization of cytosolic β-catenin, its translocation to the nucleus and downstream effects on gene transcriptional. Activation of β-catenin is also linked to regulation of dendritic remodeling. The WNT pathway in nAcc has been directly implicated in behavioral responses to CSDS ^4,44^. There is also evidence for the importance of WNT/ß-catenin signaling as a mediator of the behavioral effects of antidepressant drugs ^34^, with decreased susceptibility observed in animals exposed to CSDS upon induction of this signaling pathway in nAcc ^4,44^.

Our findings also implicated *Esr1*, for which there is evidence of coupling with the WNT-ß - catenin pathway ^45^. Estradiol acts through estrogen receptor alpha to regulate β-catenin-mediated transcription by affecting canonical WNT signalling. This effect is considered a candidate mechanism for the neuroprotective effects of estrogens ^45^. Increased estrogen signaling associates with enhanced WNT-dependent function, most notably neurogenesis and synaptic plasticity, in both male and female rodents ^46,47^. Estrogen administration produces an anxiolytic-like effect also apparent in both male and female rodents ^47–49^. Our previous studies ^16^ identify the nAcc as a critical site for such effects. Selective overexpression of *Esr1* in the nAcc enhances resilience to CSDS.

Finally, we used a recent GWAS of neuroticism to examine the convergence with the molecular pathways identified in the mouse CSDS model. Previous transcriptomic analyses as well as the current epigenomic study identified the WNT-ß-catenin pathway as a critical candidate mechanism for susceptibility to chronic stress ^4,44^. We found significant overlap between our gene list defined by DM regions in gene bodies and the top genes identified in the GWAS of human neuroticism ^3^. Enrichment analysis of the overlapping genes once again revealed the WNT-ß-catenin pathway as the top biological process (Fig.4, Fig. S1a, b). This striking level of convergence suggests a shared, candidate biological pathway linked to neurogenesis and synaptic plasticity. The findings underscore the relevance of model systems that focus on clinically-relevant phenotypes for our understanding of the molecular pathways associated with vulnerability for mood disorders.

## Methods

### Animals and behavior

All mice were obtained from Jackson Labs and maintained on a 12 hour light-dark cycle with lights on at 7:00 am and a controlled temperature range of 22-25°C. Food and water were provided *ad libitum*. All experiments were approved by the Institutional Animal Care and Use Committee (IACUC) guidelines at Mount Sinai. All behavioral testing was counterbalanced across experimental groups, and assignment to experimental groups was random. CSDS and social interaction tests were performed according to our established protocol^7,50^. Briefly, 8-week-old male C57BL/6J mice were subjected to ten daily defeats by a novel CD1 retired male breeder aggressor mouse that had been previously screened for aggressive behavior. CD1 mice were placed on one side of a large hamster cage separated by a perforated Plexiglas divider. At the onset of CSDS, a single C57BL/6J test mouse was placed into the same side of the cage as the CD1, and the CD1 was given 10 minutes to physically attack the C57BL/6J mouse. At the end of the 10 minutes, the test mouse was moved to the other side of the Plexiglass divider, where there was no more physical contact. However, as the divider was perforated, sensory contact continued. After 24 hours of sensory contact, the test mouse was moved into another hamster cage where it was defeated by a different CD1 mouse and so for 10 days. Control mice were housed in a mouse cage with a Plexiglas divider, with a novel C57BL/6J mouse placed on the opposite side of the divider, but no physical contact was allowed between the two mice. Control C57BL/6J mice were moved to a new half of a cage every day during the defeat procedure. After the final defeat, test mice were single-housed in preparation and screened in the social interaction test the subsequent day. Social interaction testing was performed under red light in a closed behavioral chamber. C57BL/6J mice were placed into an open arena with an empty cage at one side (interaction zone). Mice were given 2.5 minutes to explore the arena and then removed. A novel CD1 aggressor to which the C57BL/6J mouse had never been exposed was placed in the cage (interaction zone) and the procedure was repeated. Time in the interaction zone was recorded automatically with video tracking software. Data were analyzed as time spent in the interaction zone when the aggressor was absent compared to time spent in the interaction zone when the aggressor was present. Social Interaction ratio (SI) was calculated as 100 x [(time in the interaction zone with a target mouse present)/(time in the interaction zone with target absent)]. Susceptible mice are defined as an SI < < 0.8. This measure of susceptible predicts several other behavioral abnormalities exhibited by stress-susceptible defeated mice ^50^. Susceptible and control mice were euthanized by decapitation, and nAcc was dissected rapidly as 14 gauge punches and frozen and stored at −80°C for further analyses by WGBS.

### DNA extraction and MethylC-Seq

Genomic DNA was extracted from frozen nAcc punches using Qiagen DNA micro kit (Qiagen, Cat# 56304.). On-column RNaseA treatment was performed during DNA extraction to reduce RNA contamination.

Methylated and unmethylated DNA sets (Cat# D5017, pUC19 DNA set, Zymo research) were added as spike-in controls (2 ng spike-in control in 1 µg DNA) to each DNA sample to evaluate the bisulfite conversion efficiency. DNA samples were sheared and subjected to bisulfite conversion. The whole genome bisulfite sequencing (WGBS) libraries were prepared using Pico Methyl-seq library prep kit (Cat# D5456, Zymo research). Paired-end, 150 bp read-length DNA-seq was processed in Illumina HiSeq X at a sequencing depth of 30x genome coverage.

### DNA methylation data analysis

The quality of raw sequencing data was assessed with the FastQC program (https://www.bioinformatics.babraham.ac.uk/projects/fastqc/). Informed by this step, Cutadapt (http://cutadapt.readthedocs.io/en/stable/index.html) was used to remove the first 8 bases of reads and full or partial Illumina TruSeq adapters. Subsequent reads with fewer than 20 bases were discarded. All sequence reads were aligned to the original and bisulfite-converted mouse mm10 genome with the Bismark suite ^51^, using Bowtie1 and the default parameters which allow up to 2 mismatches within the 50 base pair seed region. BAM files from the Bismark alignment were merged and sorted with Samtools ^52^. Methylation calling and statistics were computed with methylKit ^53^. Differentially methylated bases were visualized in the mm10 genome with the Integrative Genomics Viewer (IGV) ^54^. Network and gene ontology analysis were computed with Metacore^®^ (Clarivate Analytics). In the place of fold change values from differential expression analyses usually used by this program we used as input a weighted methylation metric defined as the product of the number of DM bases in a gene and the sum its Δβ values divided by gene length. Statistical analyses were performed in R version 3.5.2. The GeneOverlap Bioconductor package was used for the significance of overlap of gene lists, and the two sided prop.test function for the difference in population proportions.

## Data Availability

Raw sequencing read files and methylation call files have been submitted to the Gene Expression Omnibus (GEO) under accession GSE126955.

## Authors’ contributions

NO performed the WGBS data analysis and wrote the paper. TYZ supervised the WGBS project, performed Clarivate Analytics MetaCore analysis and wrote the paper. XW performed DNA extraction. JD supervised part of the project. PPS analyzed part of human GWAS data with current animal data. BL performed the behavioral testing and extracted the tissue samples. EJN supervised the project and revised the paper. MJM supervised the project and together with NO wrote the initial draft paper. All authors read and agreed with the final manuscript.

## Acknowledgements

This research was funded by grants from the Hope for Depression Research Foundation (EJN and MM) and the National Institute of Mental Health (EJN).

